# Effect of combined Respiratory Muscle Training on Sleep and Cardiovascular Biometrics in a non-clinical cohort

**DOI:** 10.1101/2025.06.27.661934

**Authors:** Travis Anderson, Kerry Martin, Nina Bausek

## Abstract

**Study objectives:** Sleep disruption is a growing health problem, affecting a significant portion of the adult population worldwide. Insufficient sleep has various short- and long-term consequences, including an elevated risk of cardiovascular and metabolic diseases. Previous research has demonstrated that resistive respiratory muscle training (RMT) can enhance both sleep quality and cardiovascular health in individuals diagnosed with obstructive sleep apnea, underscoring its efficacy as a non-pharmacological therapeutic strategy for this patient group. However, the effects of RMT on sleep and cardiovascular parameters have not been investigated in non-clinical populations.

**Methods:** This prospective study investigated the effects of combined inspiratory and expiratory RMT (cRMT) on sleep parameters and cardiovascular biometrics, specifically heart rate variability (HRV), in a non-clinical adult cohort. Utilizing a wearable device for remote data collection, this randomized controlled trial included 67 participants divided into good and poor sleeper groups based on historical sleep data. During a five-week intervention period, participants in the intervention group underwent cRMT using a Breather Fit device, while control group participants did not receive the intervention.

**Results:** Study findings demonstrate a significant increase in overnight HRV metrics during the intervention period compared to the baseline, indicating improved autonomic cardiac function. However, no significant changes were observed in any parameters of sleep quality.

**Conclusion:** These results suggest that cRMT may enhance cardiovascular health by improving autonomic function in non-clinical populations without directly affecting sleep quality. This study underscores the potential of RMT as a non-pharmacological intervention to improve cardiovascular health, warranting further investigation in future studies.

**Brief summary:**

a. **Current Knowledge/Study Rationale:** Insufficient, disrupted, or ineffective sleep is prevalent and can increase the risk of developing cardiovascular disease (CVD). While respiratory muscle training has been found effective in improving sleep and cardiovascular parameters in sleep apnea, its effect in healthy people with or without sleep issues is unknown.
b. **Study Impact:** This study demonstrates significant benefits of RMT on CVD metrics including HR and HRV, indicating improved autonomic cardiac function. These findings highlight the potential of RMT as a non-pharmacological intervention to improve cardiovascular health.

## Introduction

Sleep disruption is an increasing health problem, affecting 20-35% of adults worldwide ^1^. Insufficient sleep has short and long-term consequences, including increased risk of cardiovascular and metabolic diseases, cognitive impairment, premature mortality, and reduced quality of life ^2^. In particular, sleep-breathing disorders are a major concern since they have been associated with the most common causes of death, like heart failure, myocardial infarction, and stroke ^3–6^. A recent systematic review revealed an overall hazard ratio of 1.78 (95% CI 1.34–2.35) for cardiovascular pathology in the adult population suffering from sleep-breathing disorders. Moreover, sleep-breathing disorders such as obstructive sleep apnea (OSA) increase the incidence of major adverse events in patients with cardiovascular disease ^7^. The pathophysiological mechanisms leading to cardiovascular disease in patients with OSA have been extensively studied. Hypoxemia and hypercapnia resulting from apnea episodes lead to the activation of the sympathetic nervous system, which increases peripheral vasoconstriction with consequent hypertension ^8^.

Given the intricate relationship between sleep-breathing disorders and cardiovascular health, heart rate variability (HRV) emerges as a critical non-invasive biomarker for assessing the autonomic nervous system’s response to these conditions ^9^. HRV is the variation or irregularity in duration of beat-to-beat or interbeat intervals ^10^. Interbeat intervals at rest are primarily regulated by the parasympathetic branch of the autonomic nervous system (ANS) ^10^, and thus HRV metrics serve as indicators of cardiovascular ANS function ^11^. The heightened sympathetic drive is associated with reduced HRV, while parasympathetic dominance is associated with increased HRV. In patients with OSA, the increased sympathetic tone persists during quiet wakefulness, likely due to enhanced chemoreflex gain ^8^, which is accompanied by decreased HRV ^12^. Furthermore, other sleep disruptions such as difficulty falling asleep, sleep continuity disturbance, nonrestorative sleep, and short sleep duration also increase the risk for cardiovascular disease in healthy people by increasing sympathetic tone ^2^. Lower HRV is associated with a higher risk of cardiovascular events and, in patients with acute myocardial infarction, with an increased risk of mortality ^13–16^. Furthermore, lower HRV in normotensive men has been associated with a higher risk of developing hypertension ^17^. Therefore, HRV assessment is highly clinically relevant, especially in patients with a high risk of developing cardiovascular disease or its severe complications.

Several non-invasive respiratory interventions have been shown to improve both sleep breathing disorders and HRV. Daytime breathing retraining, such as diaphragmatic breathing, singing exercises, aerobic exercises, and inspiratory muscle training, improves the apnea-hypopnea index and reduces daytime sleepiness in patients with OSA, improves sleep parameters, and reduces snoring ^18^. Respiratory muscle training (RMT) is a technique designed to enhance the functioning of the respiratory muscles through targeted exercises, increasing the workload of respiratory muscles by training against resistance. Resistive RMT requires individuals to inhale and exhale using a device with an orifice with a variable diameter, increasing the load with decreasing diameter size ^19^. This load variation creates a training effect that challenges and strengthens the respiratory muscles ^19–21^.

Resistive RMT has previously been shown to improve both sleep and cardiovascular function in patients with OSA ^22,23^, highlighting its effectiveness as a non-pharmacological and non-invasive therapeutic approach for this population. However, the effects of RMT on sleep restoration and cardiovascular autonomic function have not been evaluated simultaneously in the non-clinical population. In this proof-of-concept prospective study, we assessed sleep parameters and cardiovascular biometrics after RMT intervention by remotely recording data with a wearable device during the natural night resting period in a cohort of adult non-clinical individuals. This pilot study aimed to validate the remote assessment of sleep and cardiovascular parameters and the use of RMT to improve autonomic cardiac function in non-clinical populations, addressing the potential use case for RMT to improve sleep and autonomic balance in an apparently healthy population.

## Methods

Participants (n = 67) were selected from existing users of the Biostrap EVO (Biostrap LLC) wearable device. Participants were selected into two sub-samples based on their historical sleep characteristics. Participants who met the following criteria based on historical Biostrap data were considered good sleepers (n = 44): less than five awakenings per night, an average sleep score >70 out of 100, and an average sleep SpO_2_ of ≥96 ^24^. The sleep score was determined from factors including total sleep duration, minutes of deep sleep (estimated), number of involuntary awakenings, relative HR compared to an individual’s rolling 30 day average, and absolute number of low SpO2 readings with bins of 90-95, 80 to 89, and below 80 representing specific penalties. This gives a gross sleep score that is then corrected based on sleep efficiency, calculated as the total time asleep relative to the time in bed with a 5% buffer (no penalty). The remaining participants who did not meet these criteria were considered poor sleepers (n = 23). Within each subsample, participants were randomized into control and intervention groups. All participants signed a written informed consent form prior to participating in the study.

All participants completed a one-week baseline phase, during which participants were required to wear their wrist-worn Biostrap wearable device each night while sleeping. Following the baseline phase, all participants completed a five-week experimental phase including combined inspiratory and expiratory combined RMT (cRMT). Participants randomized to the intervention group (n = 29) were required to use the Breather Fit (PN Medical, FL, USA), an cRMT device. The device provides adjustable resistance during inhale and exhale for strengthening the inspiratory and expiratory muscle groups. The training protocol included three sets of 10 breaths, twice per day on six days of the week. The target intensity of cRMT was 70% of maximum effort and those in the intervention group were supplied with an instructional video on cRMT and device care. Participants of the control group (n = 38) received no respiratory exercise. All participants were required to continue wearing the Biostrap EVO device throughout the experimental phase. Lastly, all participants completed a one-week washout phase, during which they continued wearing the Biostrap device but ceased using the RMT device.

### Sleep Parameters and Biometrics

The Biostrap EVO device uses photoplethysmography (PPG) to measure a range of cardiovascular biometrics. Additionally, PPG and accelerometer signals are used in proprietary algorithms to assess several sleep quantity and quality metrics ^25–27^. All sleep parameters and biometrics assessed in the present study and their descriptions are presented in Table 1.

**Table 1.** Biostrap sleep and biometric dependent variables, abbreviations, and descriptions.

| Variables | Abbreviation | Description |
| --- | --- | --- |
| Sleep duration | Sleep<sub>DUR</sub> | The time between sleep onset and sleep offset (mins) |
| Sleep efficiency | Sleep<sub>EFF</sub> | Time asleep relative to total time in bed. This accounts for sleep latency and any awakenings throughout the sleep session. |
| Light sleep | Sleep<sub>LIGHT</sub> | Time in light sleep for the sleep session (mins) |
| Deep sleep | Sleep<sub>DEEP</sub> | Time in deep sleep for the sleep session (mins) |
| Arousal count | Arousal | Number of large accelerometer movements during the night |
| Awake time | Awake | Time spent awake during the sleep session (mins) |
| Sleep latency | Sleep<sub>LAT</sub> | Time from going to bed and sleep onset (mins) |
| Sleep score | Sleep<sub>SCORE</sub> | The Biostrap proprietary score for a sleep session (0-100) |
| Biometric | Abbreviation | Description |
| Heart rate | HR | The average frequency of ventricular contractions (bpm) |
| Resting heart rate | HR<sub>REST</sub> | The average of the five lowest HRs after excluding the lowest three HRs (bpm) |
| Heart rate variability | HRV | The average RMSSD over sleep session (ms) |
| Resting heart rate variability | HRV<sub>REST</sub> | linear regression projected waking RMSSD from overnight trend (ms) |
| High frequency power | HF | High frequency HRV power (AU, after Fourier transform) |
| Low frequency power | LF | Low frequency HRV power (AU, after Fourier transform) |
| Oxygen saturation | SpO<sub>2</sub> | The average oxygen saturation during a sleep session |
| Ventilation rate | Vt | The average respiratory rate during a sleep session |

Abbreviations: RMSSD = root mean square of successive intervals; bpm = beats per minute; AU = arbitrary units; ms = milliseconds; HR = heart rate.

### Analytical Approach and Statistical Analysis

Sleep and biometric data were exported from Biostrap. Linear mixed-effects models with random intercepts were used to assess the interaction between the study phase and the study group in predicting the dependent variable (i.e., sleep or biometric variables). First, data were modeled with all participants, regardless of sleep group assignment. Then, in separate models, changes in sleep parameters and biometrics were assessed using observations from only poor sleepers or only good sleepers. All models were fit using the *lmer* package^1^, and all statistical analyses were completed using R statistical software (v4.1.1).

## Results

A total of 2342 individual observations from 67 participants were included in the analyses. All models converged with random intercepts and interaction coefficients (95%CI) are reported in Table 2. Comparative models including all participants demonstrated a significant effect (significant changes over time) of the intervention on HRV, HF, and HRV_REST_ (Table 2, Figure 1), where all variables increased in the experimental phase compared to the baseline phase for the intervention group, but not the control group, with no changes in the washout phase. When restricting the analysis to good sleepers only, HRV and HRV_REST_ again demonstrated an experimental phase by intervention group interaction, wherein both biomarkers increased in the intervention phase. Furthermore, HR_REST_ significantly increased in the washout period compared to the baseline phase for the intervention group, suggesting a delayed effect on this biometric.

**Table 2.**
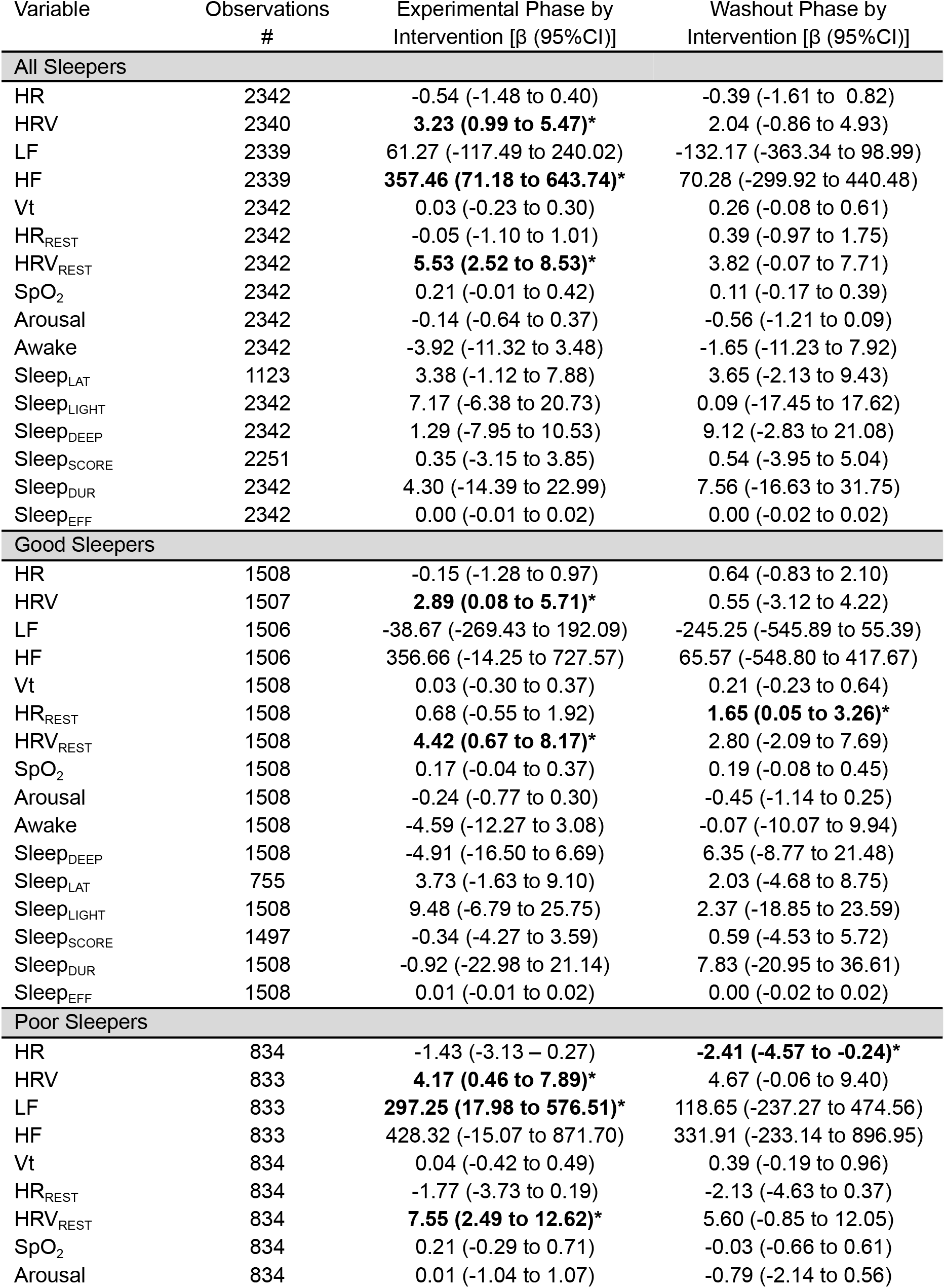
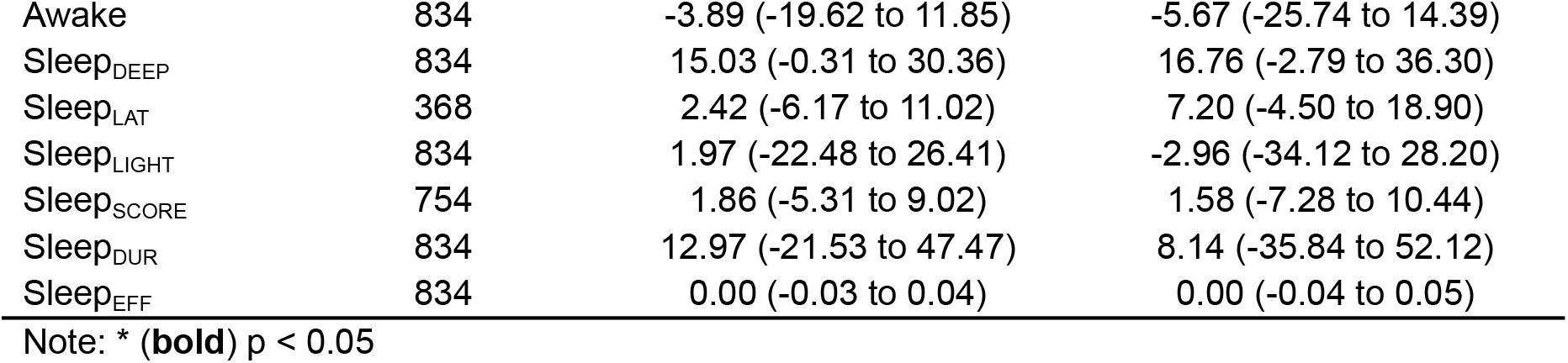
Model coefficients (95%CI) for interaction terms for all models for all sleepers, good sleepers, and poor sleepers.

**Figure 1.**
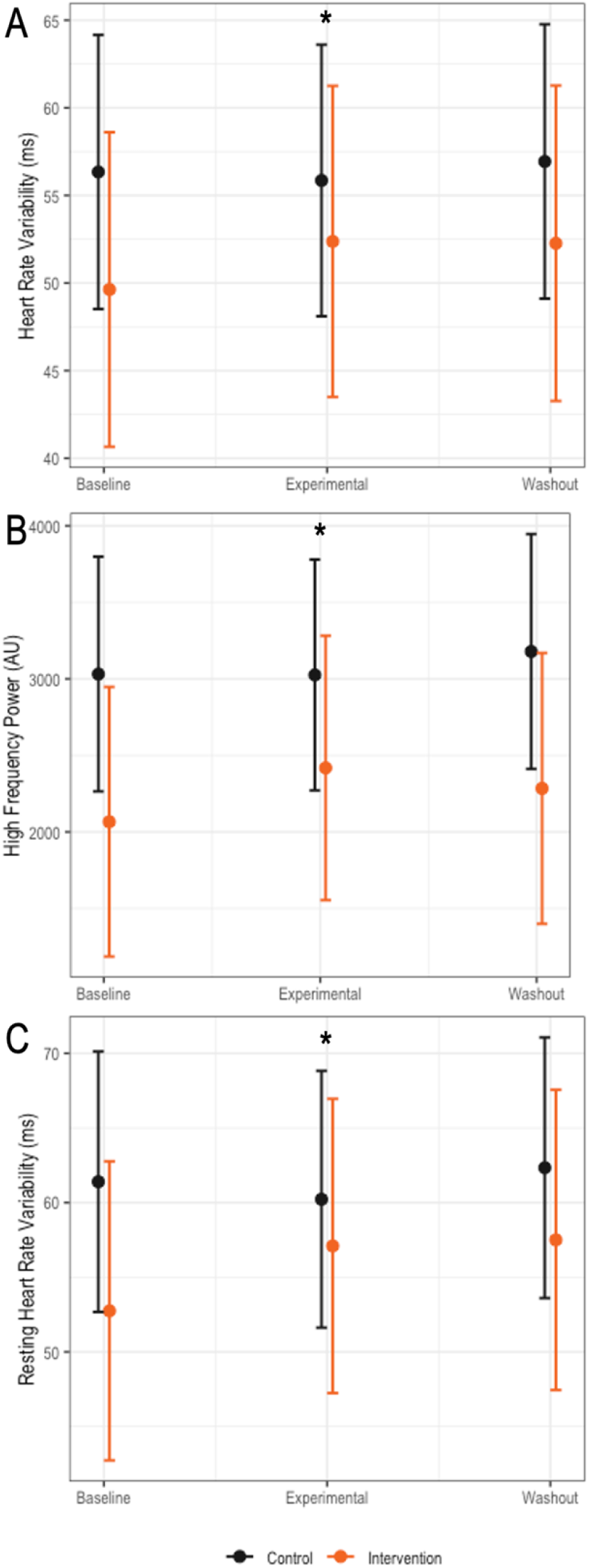
Differences in HRV between control group and total participants. Model estimates of change in HRV (A), HF (B), and HRV_REST_ (C) across study phases for all sleepers. * Indicates a significant phase by intervention interaction.

When restricting to poor sleepers only, HRV and HRV_REST_ again demonstrated an experimental phase by intervention group interaction, wherein both biomarkers increased in the experimental phase. Further, LF increased in the experimental phase for the intervention group. Lastly, HR significantly decreased in the washout period compared to the baseline phase for the intervention group. Interestingly, a non-significant increase in Sleep_DUR_ was seen in poor sleepers, but not in good sleepers (Table 2).

Comparative analysis revealed that significant phase by intervention interactions for HRV, HRV_REST_, and LF observed in poor sleepers differed from the control group during the experimental phase, but not during baseline or washout (Figure 2).

**Figure 2.**
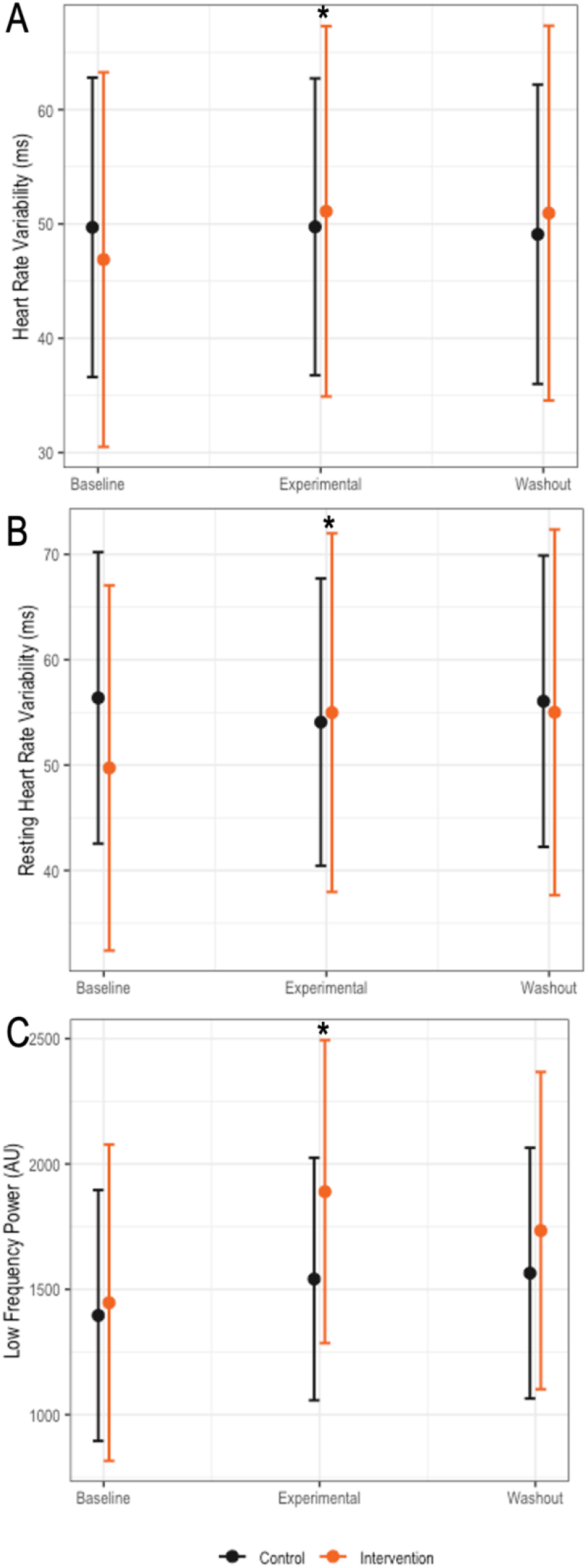
Differences in HRV between control group and poor sleepers. Model estimates of change in HRV (A), HRV_REST_ (B), and LF (C) across study phases between intervention groups for poor sleepers only. *Indicates a significant phase by intervention interaction.

## Discussion

This study shows the effects of 4 weeks of combined RMT on sleep and cardiovascular parameters in non-clinical adults with different sleeping patterns. By using a novel approach consisting of monitoring biomarkers of interest in a completely remote way, we collected and analyzed a large number of individual observations. Through subsequent data analysis, we found that RMT as the sole intervention significantly increased HRV during the study period, indicating increased activity of the parasympathetic nervous system (PNS). The intervention also decreased HR during the follow-up period, suggesting sustained benefits of autonomic improvement.

The classical framework supporting HRV as an indicator of the autonomic function states that the high frequency (HF, 0.15-0.4 Hz) component of HRV reflects the parasympathetic activity, whereas the low frequency (LF, 0.04-0.15 Hz) component may reflect altered baroreflex function and cardiac autonomic outflow, rather than sympathetic tone ^28^. In our study, RMT not only significantly increased HRV in all groups but also HF in all-sleeper combined data, suggesting that the intervention increases parasympathetic activity. Lower HRV is associated with a higher risk of cardiovascular events and higher mortality risk in patients with a previous acute myocardial infarction ^13–16^. Furthermore, the Framingham Heart Study evaluated the HRV and blood pressure in 931 men and 1111 women attending a routine examination and found that HRV was significantly lower in hypertensive people. Remarkably, the 4-year follow-up of those people who were normotensive in the first visit revealed that lower HRV at baseline was a predictor of a higher risk of developing hypertension in men ^17^. Therefore, the findings of our pilot study in a non-clinical population support a correlation between RMT and improved autonomic outcomes, with potential for cardiovascular risk reduction. Further studies are needed to confirm this association and to address the underlying mechanisms.

RMT has previously been shown to improve autonomic function in people with different disorders. In patients with OSA, resistive RMT not only improves sleep quality, daytime sleepiness, and lung function^21^ but also reduces blood pressure and plasma norepinephrine levels, highlighting its benefits on the autonomic nervous system ^23^. Moreover, RMT has been shown to improve autonomic function in patients with cardiovascular, metabolic, neurological, and respiratory diseases ^29–32^. In the present study, we found that RMT increases HRV in a non-clinical population, supporting previous studies showing the improvement of cardiac autonomic function and increase in HRV in healthy people evaluated in the waking state for 10-15 minutes in laboratory settings ^20,21,33^. Here, we further extended the recordings for hours, in individuals allocated in real-life settings during the natural night resting period, which removes external/environmental confounders and increases the ecological validity of the study. Furthermore, the data were not only collected at baseline and during the intervention phase but also after a washout period, allowing the identification of long-lasting effects, like the observed decrease in HR.

A number of mechanisms have been suggested to underlie the cardiovascular benefits of RMT that might account for the increase in HRV and HF observed in the intervention group of our study. First, several studies have shown that RMT increases the maximum inspiratory pressure, which reduces inspiratory time ^20,21,33^. Normally, inspiration increases venous return, triggering the baroreflex, which produces autonomic changes regulating the heart rate, a mechanism known as respiratory sinus arrhythmia ^34^.

Hence, it has been proposed that by strengthening the respiratory musculature, RMT increases inspiratory pressure, which results in reduced inspiratory time, and ultimately increases HRV ^21,34^. Other studies showed that RMT reduces the cardiovascular sympathetic drive by reducing respiratory muscle fatigue, presumably by reducing the activity of chemosensitive afferents within the inspiratory muscles ^35,36^. Finally, there is some evidence demonstrating that several pulmonary mechanical mechanisms induce changes in HRV independently of the autonomic nervous system ^34^.

HR not only responds to fast adjustments coming through autonomic signals but also depends on local mechanisms occurring in the sinus node, such as changes in the expression of ion channels that can be a consequence of different metabolic signals ^37^. For example, trained athletes have been shown to have different expression levels of ion channels in the sinus node that mediate the persisting bradycardia across their lives ^37,38^. In our study, a similar mechanism may explain the persistence of decreased HR after the washout period in poor sleepers.

Long-term assessment of HRV during sleep has been previously shown to be a valid tool for evaluating sympathovagal balance ^39^. Classically, it relies on data extracted from the electrocardiogram (ECG) channel of the polysomnography (PSG) and it is used to evaluate the autonomic function in the context of different disorders and interventions ^40^. The development of wearable devices using photoplethysmography and accelerometer sensors revolutionized the simultaneous assessment of sleep, cardiovascular, and other health-related variables due to the non-invasive nature of data collection in freely moving subjects. Recently, by comparing data collected from the gold-standard measures of PSG and ECG, several studies validated the use of wearable devices for the assessment of timing and duration of sleep as well as HR and HRV during the resting period of healthy subjects ^41–43^. The Biostrap wristband has also been validated for the clinical assessment of cardiovascular variables and was previously used in a clinical study for sleep monitoring ^26,44^. However, to our knowledge, this is the first study simultaneously evaluating the effects of RMT intervention on sleep and HRV in a population of healthy male and female adults by remotely collecting data from wearable devices.

Although RMT has been shown to improve sleep quality in different pathological conditions ^45–48^, in our cohort of healthy people the intervention with the RMT device has not significantly affected sleep parameters, even when restricting the analysis to those identified a priori as poor sleepers. In line with our results, another randomized study showed that RMT sustained for 4-5 weeks did not produce significant changes in sleep quality in healthy active elderly ^49^. However, we cannot exclude that a longer RMT beyond four weeks could induce significant changes in sleep parameters. In addition, we are also unable to exclude external contributing factors, such as co-sleeping or irregular working hours. It is also possible that the method for grouping caused heterogeneous outcomes.

In summary, in this proof-of-concept prospective study in which remote recordings of cardiovascular variables were performed in a cohort of healthy Biostrap adult users, combined RMT improved autonomic parameters indicative of increased parasympathetic activity but did not significantly affect sleep parameters. These findings encourage further studies performed in larger cohorts with longer time interventions to determine whether RMT is a valuable non-pharmacological strategy to improve autonomic function in healthy people as well as in different patient populations.

### Limitations

Since the study was performed remotely and on non-clinical subjects, critical medical history was not taken into account, limiting the generalizability of the findings. However, severe respiratory disorders, major surgeries or traumas, vertigo, poorly controlled hypertension, and unstable cardiac conditions were exclusion criteria assessed through a questionnaire. Moreover, we utilized a randomized allocation approach to assign participants to the intervention and control groups, which minimizes selection bias by reducing the likelihood of systematic differences between the groups at the baseline.

Another limitation pertains to the lack of sleep apnea data collection. Although the use of CPAP was an exclusion criterion, a portion of participants could have undiagnosed or untreated sleep apnea. This missing data could have implications for the study’s findings since the observed improvements in HRV could be partially attributed to the amelioration of sleep apnea rather than a direct effect of the RMT intervention. However, given that patients with sleep apnea are more likely to be poor sleepers and we found that HRV increased in both poor and good sleepers, our results suggest that RMT could have a positive impact on HRV independently of apnea improvement.

Finally, since the study was performed remotely, we did not evaluate changes in respiratory muscle strength as a result of the training intervention. However, previous studies have frequently shown that RMT at a lesser intensity than that used in this study improves maximal inspiratory pressure, a direct measure of increased respiratory muscle strength, by up to 31%^18–20^.

## Abbreviations

ANS: Autonomic nervous system
AU: Arbitrary unit
BPM: Beats per minute
cRMT: combined respiratory muscle training
CPAP: Continuous positive airway pressure
CVD: Cardiovascular disease
ECG: Echocardiogram
HF: High frequency
HR: Heart rate
HRV: Heart rate variability
Hz: Hertz
LF: Low frequency
ms: milliseconds
OSA: Obstructive sleep apnea
PNS: Parasympathetic nervous system
PPG: Photoplethysmography
PSG: Polysomnography
RMSSD: Root mean square of successive intervals
RMT: Respiratory muscle training
SpO2: Oxygen saturation
Vt: Ventilation rate

## Acknowledgments

We thank Viviana F Bumaschny for her significant contributions to valuable scientific discussions and for writing a substantial portion of the manuscript.

